# Evolution of Portulacineae marked by gene tree conflict and gene family expansion associated with adaptation to harsh environments

**DOI:** 10.1101/294546

**Authors:** Ning Wang, Ya Yang, Michael J. Moore, Samuel F. Brockington, Joseph F. Walker, Joseph W Brown, Bin Liang, Tao Feng, Caroline Edwards, Jessica Mikenas, Julia Olivieri, Vera Hutchison, Alfonso Timoneda, Tommy Stoughton, Raúl Puente, Lucas C. Majure, Urs Eggli, Stephen A. Smith

**Affiliations:** Department of Ecology & Evolutionary Biology, University of Michigan, 830 North University Avenue, Ann Arbor, MI 48109-1048, USA; Department of Plant and Microbial Biology, University of Minnesota-Twin Cities. 1445 Gortner Avenue, St. Paul, MN 55108 USA; Department of Biology, Oberlin College, Science Center K111, 119 Woodland St., Oberlin, Ohio 44074 USA; Department of Plant Sciences, University of Cambridge, Cambridge CB2 3EA, United Kingdom; Department of Animal and Plant Sciences, University of Sheffield, Western Bank, Sheffield S10 2TN, United Kingdom; Center for the Environment, MSC 63, Plymouth State University, 17 High Street Plymouth, NH 03264 USA; Department of Research, Conservation and Collections, Desert Botanical Garden, 1201 N. Galvin Pkwy, Phoenix, AZ 85008 USA; Florida Museum of Natural History, University of Florida, Gainesville, FL 32611 USA; Sukkulenten-Sammlung Zürich, Mythenquai 88, CH-8002 Zürich, Switzerland

**Keywords:** gene tree conflict, gene duplication, molecular evolution, paleopolyploidy, Portulacineae, stress adaptation

## Abstract

Several plant lineages have evolved adaptations that allow survival in extreme and harsh environments including many within the plant clade Portulacineae (Caryophyllales) such as the Cactaceae, Didiereaceae of Madagascar, and high altitude Montiaceae. Here, using newly generated transcriptomic data, we reconstructed the phylogeny of Portulacineae and examine potential correlates between molecular evolution within this clade and adaptation to harsh environments. Our phylogenetic results were largely congruent with previous analyses, but we identified several early diverging nodes characterized by extensive gene tree conflict. For particularly contentious nodes, we presented detailed information about the phylogenetic signal for alternative relationships. We also analyzed the frequency of gene duplications, confirmed previously identified whole genome duplications (WGD), and identified a previously unidentified WGD event within the Didiereaceae. We found that the WGD events were typically associated with shifts in climatic niche and did not find a direct association with WGDs and diversification rate shifts. Diversification shifts occurred within the Portulacaceae, Cactaceae, and Anacampserotaceae, and while these did not experience WGDs, the Cactaceae experienced extensive gene duplications. We examined gene family expansion and molecular evolutionary patterns with a focus on genes associated with environmental stress responses and found evidence for significant gene family expansion in genes with stress adaptation and clades found in extreme environments. These results provide important directions for further and deeper examination of the potential links between molecular evolutionary patterns and adaptation to harsh environments.

## Introduction

Temperature and water availability are two major ecological determinants of plant distribution and survival (e.g., Peel et al. 2007). Plants living in harsh environments such as low precipitation, extreme temperature ranges, intense sun and/or dry winds, often exhibit specialized morphological and physiological adaptations to abiotic stresses (Gibson 1996; Kreps et al. 2002). While the morphological adaptations are perhaps easier to identify, these long-term selective pressures are expected to leave genetic and genomic signatures in lineages adapted to extreme environments. Researchers have addressed these questions in several ways, as for example exploring the gene regions and functions associated with adaptation in certain species or small groups (e.g., Christin et al. 2007).

Over the last decade, genomic data have increased dramatically and provide new ways to address macroevolutionary questions. For example, transcrimptomic and genomic data sets provide the necessary means to more accurately identify whole genome duplication events (WGD, e.g., Jiao et al. 2011; Yang et al. 2015, Parks et al. 2018). WGDs have been identified in major clades throughout plants and have previously been suggested to be associated with adaptations to extreme environments (e.g., Stebbins 1971; Soltis and Soltis 2000; Brochmann et al. 2004), speciation/diversification (Stebbins 1971; Wood et al. 2009), and success at colonizing new regions (Soltis and Soltis 2000). We continue to identify more WGDs as we increase taxon sampling (Yang et al. 2018). Broadly sampled genomic data also facilitate analyses of lineage specific molecular evolution such as gene family expansion and differential selection among genes and taxa. Finally, genomic data allow for more thorough and accurate phylogenetic reconstruction. In particular, recent studies have illustrated that phylogenetic conflict is common, but the impact of this conflict on phylogenetic reconstruction varies across the tree of life due to different macroevolutionary processes (e.g., Shen et al. 2017; Walker et al. 2017). Genomic data allow us to examine conflict and identify the processes that contribute.

The Portulacineae comprise ~ 2200 species in nine families and 160 genera (Nyffeler and Eggli 2010a; Angiosperm Phylogeny Group 2016) that are inferred to have diverged from its sister group, the Molluginaceae, ~54 Mya (Arakaki et al. 2011). Species in this suborder are incredibly morphologically diverse, ranging from annual herbs to long-lived stem succulents to trees, and include several iconic plants such as the cacti and Malagasy Didiereaceae. They also exhibit variable habitat preferences ranging from rainforests to deserts (Smith et al. 2018a). While some species are distributed worldwide, the majority are restricted to seasonally dry areas of the northern and southern hemisphere, under either hot arid or cold arid conditions (Hernández-Ledesma et al. 2015). Many specialized traits such as fleshy or succulent stems, leaves, and/or underground perennating structures, have arisen in association with the adaptation of this clade to xeric, alpine and arctic environments. Portulacineae also include several transitions to Crassulacean acid metabolism (CAM) and C_4_ photosynthesis that improve photosynthetic efficiency and hence minimize water loss in hot and dry environments compared to C_3_ photosynthesis (Edwards and Walker 1983; Borland et al. 2009; Edwards and Ogburn 2012).

Several clades within the Portulacineae have adapted to a wide array of harsh environments. The Cactaceae encompass roughly 80% of species in Portulacineae (~1850 species; Nyffeler and Eggli 2010b) and represent perhaps the most spectacular radiation of succulent plants (Mauseth 2006; Arakaki et al. 2011). In addition to succulent structures that enable water storage, Cactaceae have an array of adaptations to cope with arid and semiarid conditions. For example, many have spines that reduce air flow on surface area for evapotranspiration, and deter herbivores (Anderson 2001). They also have extensive, but relatively shallow, root systems that facilitate quick absorption of rainfall (Gibson and Nobel 1990). The Didiereaceae also exhibit specialized adaptations, including thorns and succulent leaves, to the hot and seasonally dry environments of Madagascar. Unlike the Cactaceae and Didiereaceae that are typically found in tropical and subtropical areas, the Montiaceae have a more cosmopolitan distribution with most species occurring in colder environments. Some species of Montiaceae, especially in the genera *Claytonia* and *Montia,* are adapted to high alpine zones and/or the high Arctic (Ogburn and Edwards 2015; Stoughton et al. 2017). Repeated evolution into harsh environments suggests the possibility that the clade as a whole may be predisposed to adaptation to extreme environments.

Recent phylogenetic work has helped to resolve many relationships within the Portulacineae, but the earliest diverging relationships are only moderately or poorly supported (Arakaki et al. 2011; Ogburn and Edwards 2015; Moore et al. 2017; Yang et al. 2018; Walker et al. 2018a). This may reflect a rapid radiation, uncertainty due to gene tree conflict or whole genome duplications, or other macroevolutionary processes. Recent studies have inferred several whole genome duplication (WGD) events (i.e., paleopolyploidy) within the Portulacineae, including one at the base of the clade (Yang et al. 2018). However, Arakaki et al. (2011) primarily used chloroplast loci and Yang et al. (2018) employed transcriptomes, but taxon sampling within Portulacineae was limited. Increased transcriptomic sampling may allow for the detection of additional WGD events.

In addition to phylogenetic analyses, genomic-scale data also facilitates extensive molecular evolution analyses. In particular, these data can provide preliminary, but essential, information on the genetic basis of adaptation to abiotic stresses through gene duplication and other evolutionary processes. For example, genes that have been identified to function in response to drought, cold, and heat conditions such as those involved in the abscisic acid (ABA) signaling pathway [e.g., ABRE-binding protein/ABRE-binding factors, the dehydration-responsive element binding (DREB) factors, and the NAC transcription factors (e.g., Qin et al. 2004, Nakashima et al. 2012)] provide good candidates for detecting common signatures in plant lineages, such as Portulacineae, with stress adaptation. Of course, plant stress responses can be complicated by the occurrence of multiple stresses (Nakashima et al. 2014), and how genes mediate stress responses in most plant taxa are still poorly known. Nevertheless, molecular analyses of potential stress related genes provide essential information for future targeted studies. Because the Portulacineae include many species that occupy high stress environments, they represent an excellent clade with which to examine the genetic basis of adaptation to abiotic stresses. By examining transcriptomes, we can identify lineage-specific expansions and/or positive selection among taxa that occupy these environments and gain new insights into the evolution of stress responses that may be broadly applicable to plants.

In this study, we analyzed 82 transcriptomes, 47 of which were newly generated, to better understand the evolutionary history of the Portulacineae. By closely examining patterns of gene duplication and paleopolyploidy, gene-tree/species-tree conflict, and clade-specific evolutionary patterns of genes associated with stress-related responses, we aimed to 1) identify additional WGDs; 2) examine gene tree conflict as a source of uncertainty in the reconstruction of relationships within the Portulacineae; 3) estimate phylogenetic signal for particularly contentious relationships; and 4) examine whether gene/genome duplications were associated with clade-specific adaptive traits and/or plant diversification rate.

## Results and Discussion

### Phylogeny of Portulacineae

Our concatenated supermatrix contained 841 gene regions and had a total aligned length of 1,239,871 bp. Gene and character occupancy were 95.4% and 84.5%, respectively. Of the ingroup taxa, only *Pereskia aculeata* had a gene occupancy less than 80%, while the majority (67 taxa) had over 90% of genes present in the final supermatrix. Both the concatenated maximum likelihood tree (hereafter the CML tree) and the maximum quartet support species tree (MQSST) recovered the same topology with similar support values (fig. 1), all of which were largely consistent with previous estimates of the Portulacineae phylogeny (e.g., Arakaki et al. 2011; Moore et al. 2017) despite different datasets and datatypes. For example, our analysis recovered the ACPT clade (Anacampserotaceae, Cactaceae, Portulacaceae and Talinaceae; Nyffeler 2007; Nyffeler and Eggli 2010b; Ocampo and Columbus 2010; Arakaki et al. 2011; Moore et al. 2017), with high support across genes trees for a topology of (((A, P), C), T) (figs. 1-2) that generally agrees with that of Moore et al. (2017) using a targeted enrichment approach. However, because bootstrap values may be a poor indicator of support in large phylogenomic datasets (e.g., erroneous topologies can be increasingly supported as more sequence data are added, Alfaro et al. 2003; Phillips et al. 2004), we also conducted gene tree conflict analyses (fig. 2, Smith SA et al. 2015). For example, while we found strong support (100% bootstrap for both MQSST and CML trees) for the sister relationship of Molluginaceae and Portulacineae (fig. 1), consistent with several recent studies (Edwards and Ogburn 2012; Yang et al. 2015, 2018; Moore et al. 2017), gene tree analyses highlighted that most genes were uninformative for or conflicted with this resolution. We discuss the ACPT resolution in more detail below.

**Fig. 1.**
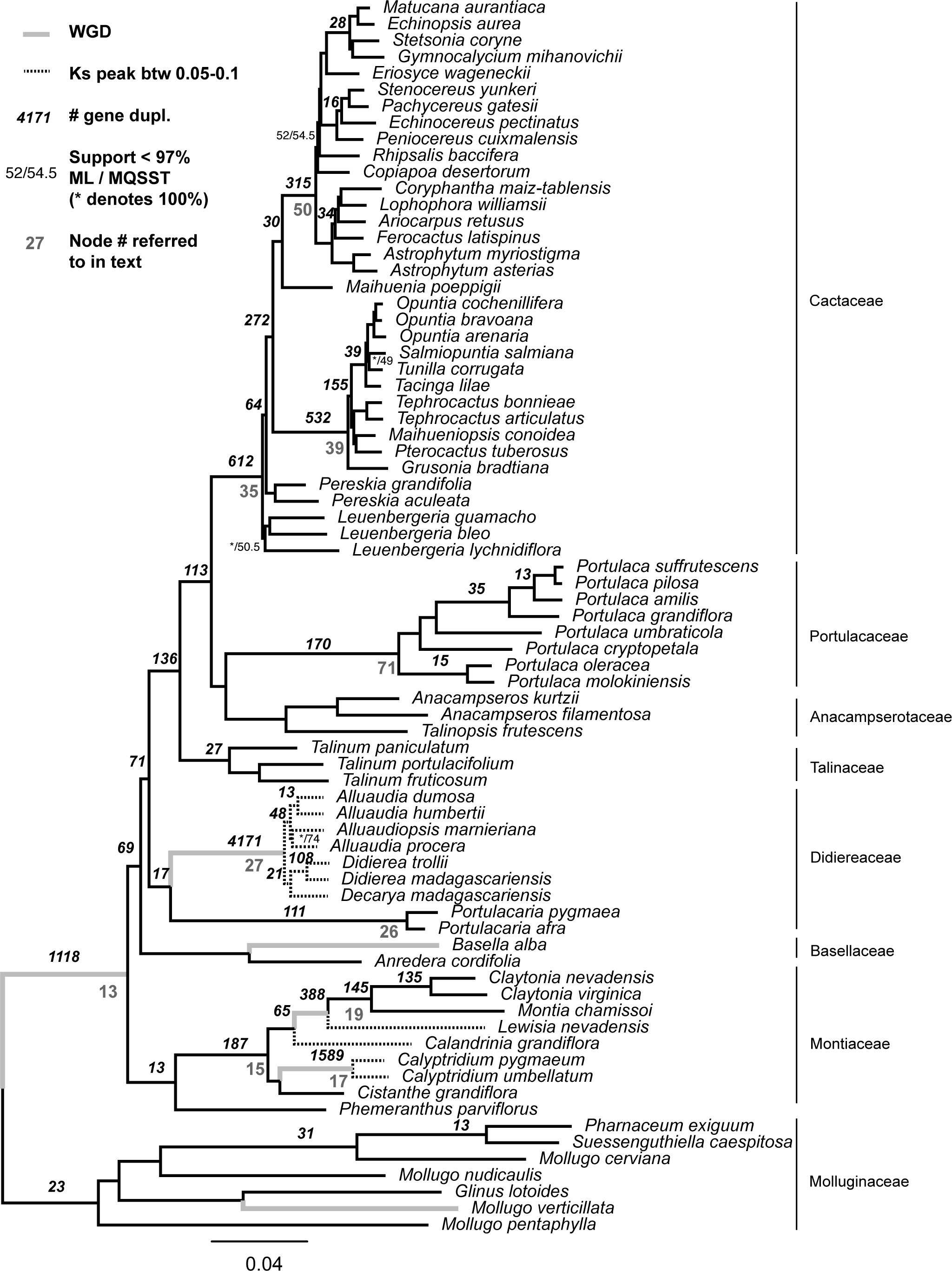
Species tree from RAxML analysis of the 841-gene supermatrix. Gene duplication numbers (i.e., the number of duplication events; number > 12 are shown) are calculated only for branches with strong support (SH-like > 80) in the 8332 rooted clusters.

**Fig. 2.**
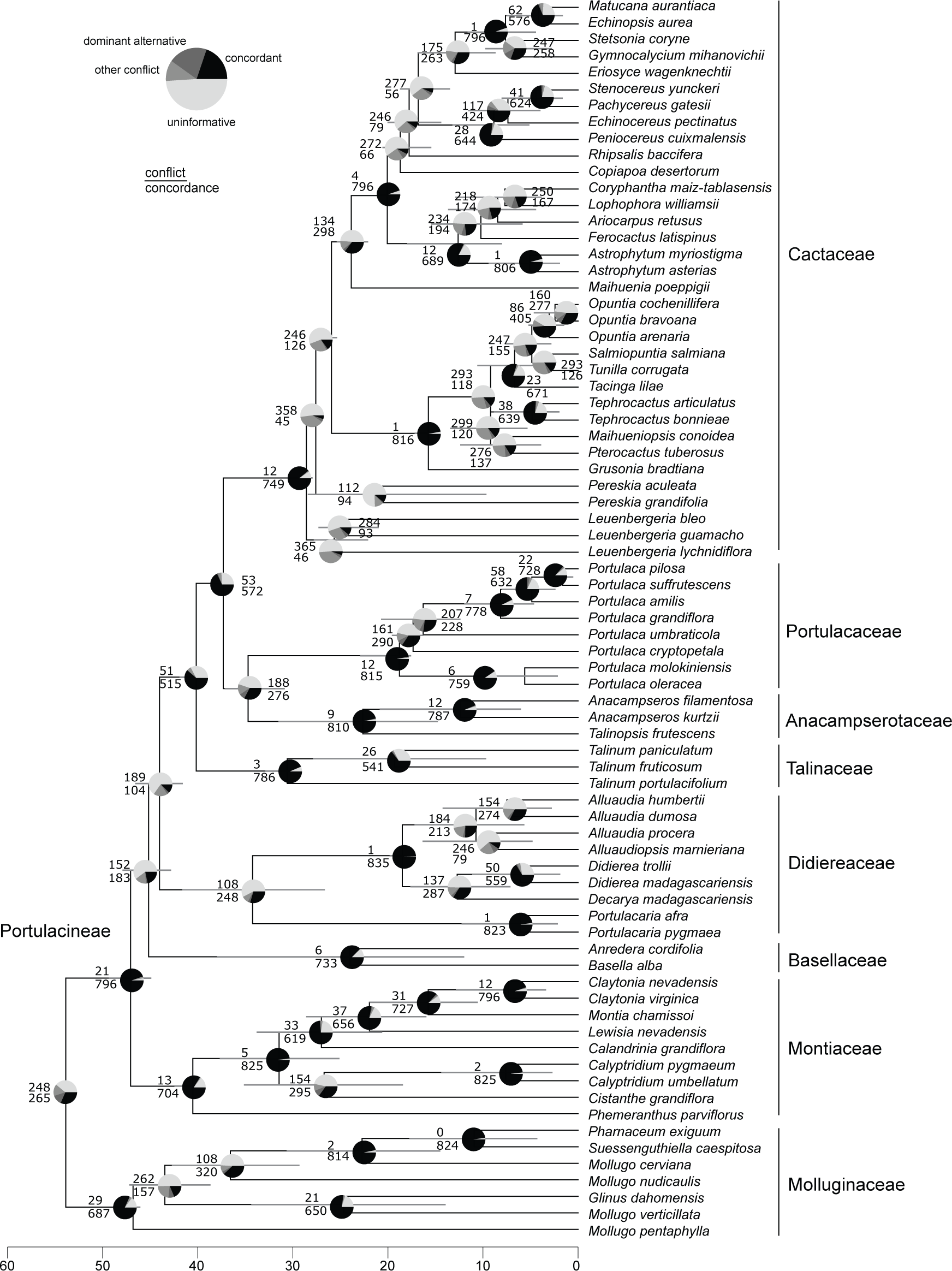
Gene-tree conflict. The pie chart proportions represent genes supporting congruent relationships (concordant), genes supporting the most common conflict (dominant alternative), all other conflicting gene trees (other conflict), and genes with no support (uninformative).

Our broad taxon sampling also allowed for resolution of relationships among early-diverging lineages within Portulacineae families. We found support for the sister relationship of Didiereaceae subfamilies Portulacarioideae (*Ceraria* and *Portulacaria*) and Didiereoideae (Madagascar Didiereaceae; Applequist and Wallace 2003; fig. 1). Although many gene trees were uninformative at this node, the ML topology was also the most frequent found among the gene trees (fig. 2). Relationships among the genera within the Didiereoideae have been difficult to resolve with targeted-gene analyses (Applequist and Wallace 2000; Nyffeler and Eggli 2010b). Here, we recovered strong support (100%) in gene tree analyses for a clade including only *Alluaudia* and *Alluaudiopsis* (fig. 1). This result differs from Bruyns et al. (2014) that found *Alluaudiopsis* to be sister to the remaining Didiereoideae.

Within Cactaceae, we recovered three major clades (*Pereskia* s.s., *Leuenbergeria,* and the core cacti, as in Edwards et al. 2005 and Bárcenas et al. 2011) with strong support (> 97%, fig. 1), but with substantial gene tree discordance (see below, fig. 2). Most genes (>85%) were uninformative or in conflict (fig. 2) among the earliest diverging nodes of Cactaceae. Within the core cacti, we recovered *Maihuenia* as sister to Cactoideae with strong support, and this clade as sister to Opuntioideae (fig. 1) as found by Edwards et al. (2005) and Moore et al. (2017). Nevertheless, the position of *Maihuenia* has been found to be highly unstable (e.g., within Opuntioideae, Butterworth and Wallace 2005; sister to Opuntioideae, Nyffeler 2002; or sister to Opuntioideae+Cactoideae, Hernández-Hernández et al. 2014; Moore et al. 2017). Unlike other areas within the Cactaceae that have high gene tree discordance, the monophyly of Opuntioideae (e.g., Barthlott and Hunt 1993; Wallace and Dickie 2002; Griffith and Porter 2009; Hernández-Hernández et al. 2011) and Cactoideae (Nyffeler 2002; Bárcenas et al. 2011) had relatively low conflict (95-97% gene trees support these monophyletic groups; fig. 2). Within Opuntioideae, however, there was a high level of discordance among gene trees. However, these conflicts may have been influenced by limited taxon sampling as we only have one to six species representing each tribe (figs. 1 and 2). The topology of the Cactoideae was largely congruent with previous studies (e.g., Butterworth et al. 2002; Nyffeler 2002; Bárcenas et al. 2011; Hernández-Hernández et al. 2011). Within Cacteae, the relationships among the five sampled genera were strongly supported and are consistent with relationships recovered by Hernández-Hernández et al. (2011) and Vázquez-Sánchez et al. (2013) using five loci. However, the core Cactoideae (sister to Cacteae, Hernández-Hernández et al. 2011) were poorly supported in both the CML tree (52%) and the MQSST inference (54.5%, fig. 1). With limited taxon sampling, some relationships (e.g., the monophyly of the Pachycereeae) are largely consistent with previous studies (e.g., Nyffeler and Eggli 2010b), whereas others (e.g., the position of *Gymnocalycium*) are still incongruent among studies (Arakaki et al. 2011; Bárcenas et al. 2011; Hernández-Hernández et al. 2014).

### Assessing conflict at specific nodes

Incongruence can result from several biological processes, including incomplete lineage sorting (ILS), hybridization, and horizontal gene transfer (HGT). As more genomes and transcriptomes have become available, our ability to analyze phylogenetic conflict and concordance across hundreds or thousands of genes has increased significantly. We found that gene tree conflicts are prevalent within Portulacineae. Several areas with high conflict occurred immediately after inferred WGD events (fig. 2) suggesting a major role of gene duplication and loss.

While incorporating discordance into species tree analyses is important (Liu and Pearl 2007; Liu et al. 2008), a close examination of gene-specific discordance as it pertains to support, or lack thereof, for different clades is also valuable. Recently, studies have also demonstrated that only a few genes in phylogenomic datasets can drive the resolution of nodes (e.g., Brown and Thomson 2017; Shen et al. 2017; Walker et al. 2018b). We examined individual genes and their support for specific phylogenetic relationships that had previously been identified to have significant conflicts including the earliest divergences of Cactaceae (particularly the relationships among *Pereskia* s.s., *Leuenbergeria*, and the core cacti; Edwards et al. 2005; Moore et al. 2017), the relationships among Portulacaceae, Cactaceae, and Anacampserotaceae, and the positions of Basellaceae and Didiereaceae (e.g., Moore et al. 2017, Walker et al. 2018a). This focused analysis allowed us to isolate the signal at particular nodes without having to accommodate the many diverse processes that may shape conflict in other parts of the tree. To address the early diverging Cactaceae, we examined the support for (*Leuenbergeria*, (*Pereskia* + core cacti)) versus ((*Leuenbergeria* + *Pereskia*), core cacti) (fig. 3). 472 genes (385 out of which have Δln *L* > 2) supported the grade, with 368 (287 out of which have Δln *L* > 2) supporting *Leuenbergeria + Pereskia*. We also found that one gene (cluster4707, with homology to the “NF-X1-type zinc finger protein”) strongly supported *Pereskia +* core cacti (>110 ln *L* units), although with no lineage-specific positive selection, gene duplication, or any obvious alignment problems. Overall, these data support the resolution of (*Leuenbergeria*, (*Pereskia* + core cacti)).

**Fig. 3.**
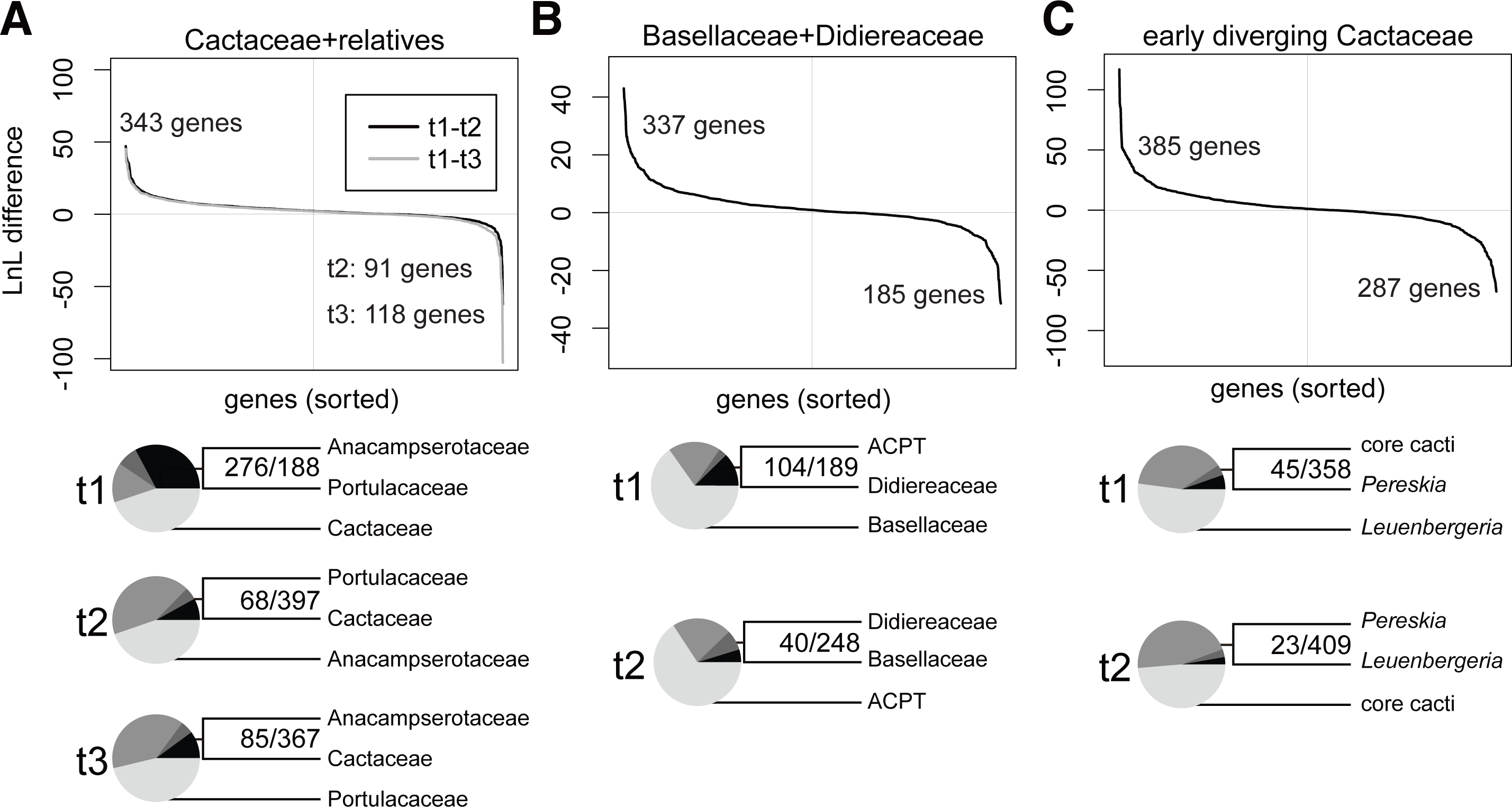
Comparison of ln *L* (sorted by greatest differences in likelihood, Δ ln*L*) for alternative phylogenetic hypotheses. Left: Three alternative resolutions were compared involving Cactaceae, Portulacaceae, and Anacampserotaceae. Values above the x-axis show the difference in likelihood for those genes supported the resolution found in our result “t1”. Those falling below the axis support “t2” (black) or “t3” (gray). Middle: Two alternative hypotheses involving Basellaceae, Didiereaceae and ACPT (Anacampserotaceae, Cactaceae, Portulacaceae, and Talinaceae). Right: Two alternative hypotheses involving the early diverging Cactaceae. Pie chart proportions are the same as in fig. 2. Outlying genes (those genes with extremely high support) are discussed in the Results and Discussion.

The relationship of Portulacaceae, Anacampserotaceae, and Cactaceae have also been contentious. Portulacaceae (P) have been recovered as sister to Anacampserotaceae (A, Ogburn and Edwards 2015; Walker et al. 2018a; and recovered here), to Cactaceae (C, Arakaki et al. 2011), and to both (Nyffeler 2007, Nyffeler and Eggli 2010). We found that 476 of 841 genes supported A+P (343 with Δln *L* > 2), 164 genes supported C+P (91 with Δln *L* > 2), and 201 genes supported C+A (118 with Δln *L* > 2). While several processes may contribute to this conflict, the distribution of the alternative placements suggest ILS to be a source of conflict (as in Moore et al. 2017) though missing data complicates our ability to test this pattern. Gene tree conflict analyses, when requiring bootstrap support > 70%, confirm this as 276 genes were congruent with the species tree, 68 were congruent with C+P, and 85 were congruent with C+A (fig. 3A). Extensive gene duplications can confound phylogenetic reconstruction due to “incomplete sorting” of paralogs (see Moore et al. 2017). However, we find low level of gene duplications at the origin of PAC (1.3%, fig. 1). We found several outlying genes including one (cluster4488, homologous to the “ARABILLO 1-like” in *Beta vulgaris*) that supported C+P (>60 ln *L* units) and one (cluster7144, homologous to the “UV-B induced protein chloroplastic-like” gene in *Chenopodium quinoa*) that supported C+A strongly (>100 ln *L* units). We did not detect any obvious errors in orthology inference (e.g., Brown and Thomson 2017), alignment, or lineage-specific positive selection.

We conducted similar analyses for the resolution of Basellaceae and Didiereaceae (fig. 3B). 506 genes (337 with Δln *L* > 2) supported (Basellaceae, (Didiereaceae+ACPT)) and 334 genes (185 with Δln *L* > 2) supported Basellaceae+Didiereaceae as found by Soltis et al. (2011) and Anton et al. (2014). No significant outlying genes (the Δln *L* ranging from -31.41 to 42.98 smoothly) were observed supporting either topology (fig. 3B).

All of the tests discussed demonstrated significant conflict among the gene regions. To address whether strong signal for hybridization contributed to the conflicting signal, we conducted hybridization analyses. The results from these analyses suggest that we cannot rule out ancient hybridization (table S1 and fig. S1). However, there is no other evidence (such as species history and geographical distribution) for hybridization between these lineages, and so we consider this to be further evidence of complex gene family evolution within the clade. Nevertheless, future analyses of synteny or other evidence may provide more direct evidence for or against hybridization as a source of conflict.

### Multiple whole-genome duplication events in Portulacineae

Whole-genome duplications (WGD) have had a profound influence on the evolutionary history of plants (e.g., Cui et al. 2006; Soltis and Soltis 2009; Jiao et al. 2011; Wendel 2015; Yang et al. 2015; Smith et al. 2018a). Previous analyses have identified WGD events within the Caryophyllales (Yang et al. 2015, 2018; Walker et al. 2017) including one in the ancestor of Portulacineae and two within Portulacineae: in the lineage of *Basella alba*, and within Montiaceae. Improved taxon sampling enabled us to confirm these events and infer additional putative WGD events as well as several instances of large-scale gene duplications (fig. 1). Ks plots have been widely applied to infer WGDs but are not without limitations (Parks et al. 2018). Higher Ks values (e.g., >2) are associated with increasingly large error (Vanneste et al. 2013) and smaller Ks values (e.g., < 0.25) can be difficult to interpret due to the errors in assembly, splice variants, or other anomalies. We used both gene tree and Ks analyses to identify WGDs more confidently (fig. 1).

We found evidence for a previously identified WGD event in the ancestor of Portulacineae (Yang et al. 2015, 2018) given the high percentage of gene duplications (13.4%) and a Ks peak (~0.4-1.0) shared by all members in this clade. However, we did not detect a similar pattern of gene duplications at the ancestor of Portulacineae+Molluginaceae (1%), confirming that Yang et al. (2018)’s estimation could result from phylogenetic uncertainty. We found a high percentage of gene duplications at the base of the Didiereoideae (50.1%) accompanied by a very recent Ks peak (between 0.05-0.10) for all the species within this clade. We note that the Didiereoideae is a relatively slow evolving lineage with a recent Ks peak. Therefore, we rely heavily on corroborating both the Ks analyses and gene tree analyses in order to confidently identify the WGD (i.e., Ks peak occurred < 0.1, fig. S2). We also identified genome duplications within the Montiaceae. Yang et al. (2018) identified WGD in the ancestor of *Claytonia* species and, with expanded sampling, we placed this WGD event at the ancestor of *Lewisia* and *Claytonia* as all four species in this clade exhibited Ks peaks between 0.2–0.4 (figs. S2 and S3). We also identified a high number of gene duplications in the two *Calyptridium* species, accompanied by a very recent Ks peak (between 0.05–0.10; fig. 1).

While we identified many gene duplications (7.3%) at the origin of the Cactaceae, this was not corroborated by Ks analyses. This is consistent with other recent studies (Walker et al. 2017; Yang et al. 2018). Several other nodes (e.g., nodes 50 and 39) exhibited similar patterns with high numbers of duplications (3.8-6.4%) but no corresponding Ks peak. It is unclear what events may be associated with these duplication events or whether gene loss, gene tree conflict, life history shifts, etc., may be obscuring the detection of gene/genome duplications, but further investigation will help shed light on these patterns and processes.

Although gene-tree-based methods can help identify the phylogenetic location of WGD events, they are unable to infer WGD events that have occurred on terminal branches. To identify these, we examined Ks peaks that are unique to one taxon and absent from close relatives. We inferred two such cases: on the branch leading to *Basella alba* (Ks=0.45) and the branch leading to *Mollugo verticillata* (Ks=0.25; fig. S2). Both species also exhibit higher chromosome numbers relative to close allies and are consistent with a previous study (Yang et al. 2018).

### WGD, diversification, and climatic niche shifts

Smith et al. (2018a) examined the potential associations between WGDs, diversification rate shifts, and climatic niche shifts within the Caryophyllales and found that some WGD events were associated with shifts in climatic niche but not diversification rate shifts. Here, we conducted similar analyses with a focus on the Portulacineae. Specifically, we examined mean annual temperature, mean annual precipitation, and the first axis of a principal component analysis on bioclimatic variables 1-19 (PCA1) among clades with WGDs (Portulacineae, Didiereoideae, and Montiaceae), as well as the Cactaceae using a species-level phylogeny. The Montiaceae, except for *Phemeranthus*, were treated together as the species-level tree conflicted with the transcriptomic tree regarding internal resolution of clades within the Montiaceae. Consistent with Smith et al. (2018a), WGD events were associated with shifts in climate tolerance (fig. 4A and 4B). Montiaceae was associated with movement into colder environments, as found in Smith et al. (2018a); Didiereoideae occupied a wetter climate than the sister clade but with both clades occupying relatively seasonally dry areas (fig. S4). The Cactaceae exhibited more variable rates of climatic niche evolution (fig. 4B), occupied a slightly hotter environment, but did not exhibit a strong pattern of different climate occupancy. This does not negate climate expansion within the clade, but suggests it may not be experienced by the entire clade.

**Fig. 4.**
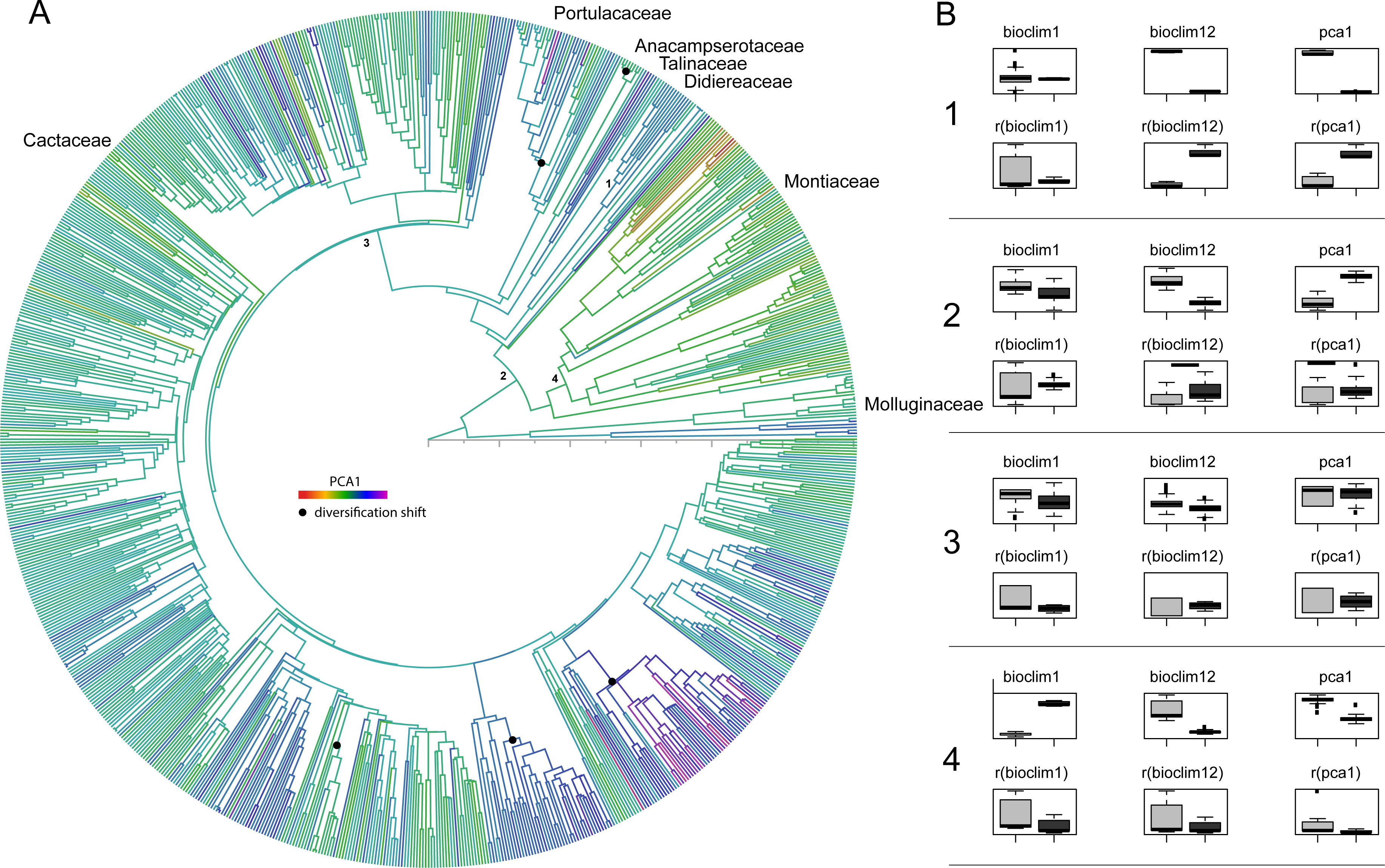
A. Reconstruction of first axis of principal component analyses (PCA1) on the species level tree of Portulacineae. Black nodes indicate the diversification rate shifts. B. Comparison of MAT (mean annual temperature), MAP (mean annual precipitation), and PCA1 between clades with WGD/extensive gene duplication (1: Didiereoideae, 2: Portulacineae, 3: Cactaceae, 4: Montiaceae except for *Phemeranthus*) and their corresponding sister clades. The upper panel is the comparison of reconstructed state. The lower panel is the comparison of evolutionary rate. Box plots are constructed from bootstrapped phylogenies.

While WGDs may not always be associated with shifts in niche occupancy, their association appears to be common within the Portulacineae and the broader Caryophyllales. The Cactaceae did not exhibit a WGD but instead extensive gene duplication and the climatic niche results for that clade are less clear. Some have suggested that WGDs may be associated with diversification rate increases (Landis et al. 2018). We examined this question on both the species-level phylogeny and a reduced phylogeny of major lineages using the estimated diversity. In the species level phylogeny, we found diversification rate shifts within Portulacaceae, Anacampserotaceae, and Cactaceae (fig. 4). In the reduced phylogeny, we found shifts at the origin of core Cactaceae (fig. S5). The differences between these are the result of the species level phylogeny being able to better place shifts while the reduced phylogeny better able to represent to total diversity. Nevertheless, in either case we did not find a correspondence between diversification shifts and WGDs.

### Gene families show broad expansions across Portulacineae

Examining gene family expansion in depth has been instrumental in understanding functional trait evolution in Caryophyllales. For example, gene expansion and neo-functionalization have been implicated in the evolution of betalain pigmentation in Caryophyllales (Yang et al. 2015; Brockington et al. 2015). Here, we explored gene family expansion in Portulacineae with respect to stress tolerance. The top 20 most expanded gene families in Portulacineae (table 1) included genes encoding transporters, proteases, cytoskeletal proteins and enzymes that are involved in photosynthetic pathways. Several genes were associated with responses to drought: the gene encoding plasma membrane intrinsic protein (PIP) can regulate the transport of water through membranes (Vandeleur et al. 2009), heat shock proteins (Kiang and Tsokos 1998), and ubiquitin (Hochstrasser 2009). But perhaps the most interesting gene expansions involved those encoding phosphoenol-pyruvate carboxylase (PEPC, Silvera et al. 2014) and NADP-dependent malic enzyme catalyses (Ferreyra et al. 2003), which are key enzymes involved in CAM/C_4_ photosynthesis (Smith and Winter 1996; Cushman 2001). Lineage-specific selection analyses on these two genes revealed that 10-13 internal branches are likely under positive selection, with the most occurred within Cactaceae. The role of gene duplication in the evolution of C_4_ photosynthesis has been contentious, and some authors have proposed that neo-functionalization of genes following duplication has not played a major role in the evolution of C_4_ photosynthesis (e.g., Williams et al. 2012; van den Bergh et al. 2014). However, recent analyses provide evidence that duplication and subsequent retention of genes are associated with the evolution of C_4_ photosynthesis (Emms et al. 2016), specifically including the NADP-dependent malic enzyme.

**Table 1.**
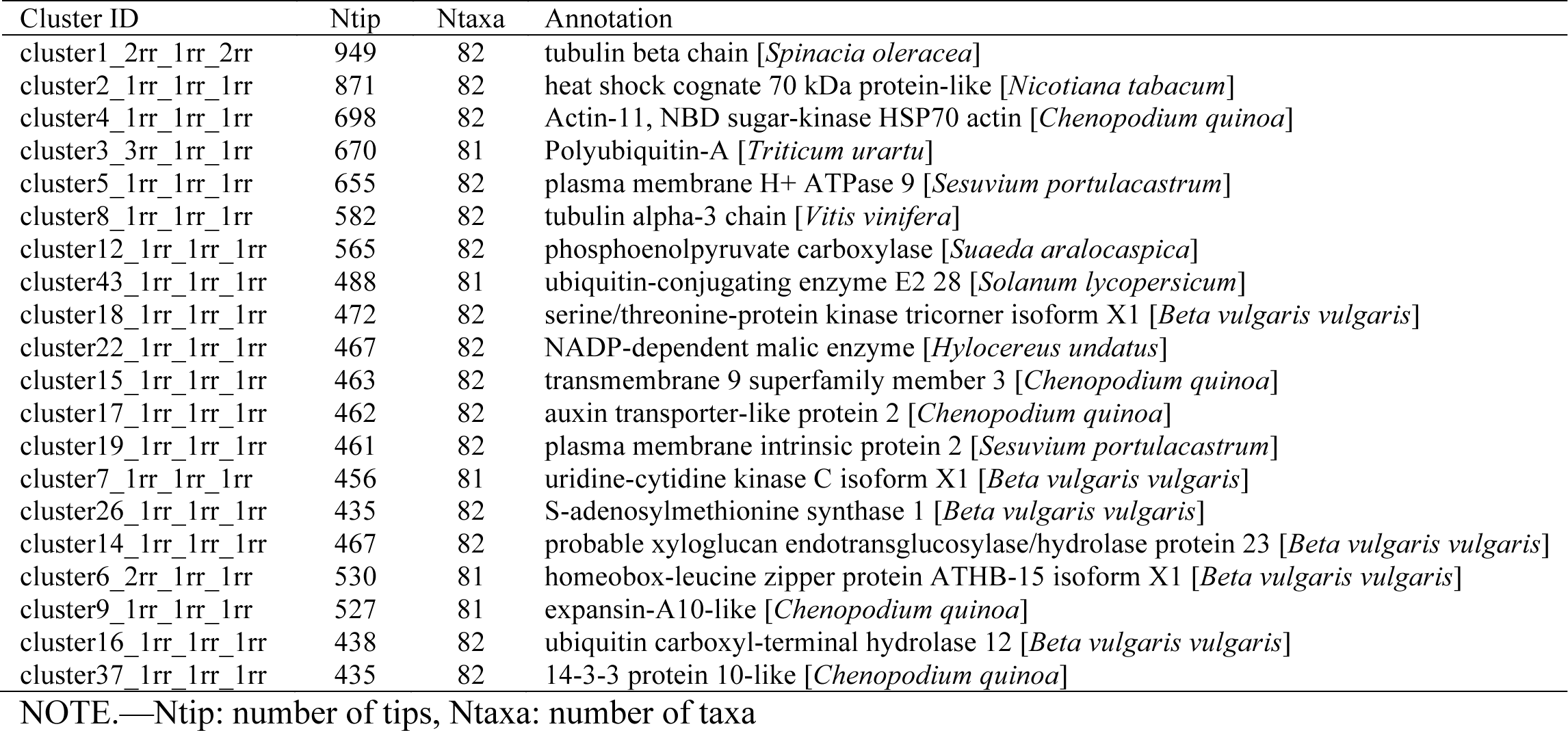
Portulacineae gene families with the highest total number of tips

Moreover, convergent evolution in several key amino acid residues of PEPC has been suggested to be associated with the origin of both C_4_ and CAM (Christin et al. 2007; Christin et al. 2014; Goolsby et al. 2018). A recent study has also confirmed the occurrence of multiple rounds of duplication within the major PEPC paralog (PEPC1E1) in the ancestral Portulacineae (Christin et al. 2014). The results presented here provide additional evidence that gene family expansion may play an important role in some aspects of photosynthetic pathway evolution.

### Lineage-specific gene expansions associated with adaptive traits

In addition to investigating genes that experienced significant expansion, we also conducted Gene Ontology (GO) overrepresentation analyses on genes duplicated at biologically significant nodes (nodes 15, 17, 19, 26, 27, 35, 39, 50, and 71; fig. 1). Of genes duplicated at each clade, 30%–55% had corresponding *Arabidopsis* locus IDs and almost all of these could be mapped in PANTHER (table S2).

We identified GO overrepresentation at three nodes (15, 27 and 35) with the Didiereoideae (node 27) exhibiting the most. This may be the result of a more recent WGD. Overrepresented GOs reflected diverse functional classes: genes belonging to “calcium ion binding” (GO:0005509) at the origins of the Montiaceae (except *Phemeranthus parviflorus*) and the Didiereoideae, genes belonging to “sulfur compound metabolic process” (GO:0006790) at the origin of Didiereoideae, and genes involved in “Hydrolase activity, acting on ester bonds” (i.e., esterase, GO:0016788, table S3) at the origin of Cactaceae. Of particular note are the genes belonging to the “sulfur compound metabolic process” (e.g., *AtAHL*, which belongs to the Arabidopsis HAL2-like gene family, Gil-Mascarell et al. 1999; and *NIFS1*, which belongs to the cysteine desulfurase family, Schmidt 2005) as sulfur-bearing evaporite compounds (principally gypsum) are commonly found in soils in areas with low rainfall and high evaporation rates (Watson 1979) where many Portulacineae lineages have diversified. Previous studies have also connected primary sulfur metabolism (e.g., sulfate transport in the vasculature, its assimilation in leaves, and the recycling of sulfur-containing compounds) with drought stress responses (e.g., Chan et al. 2013). Consequently, duplication and overrepresentation of these sulfur metabolic genes could potentially be evidence of adaptation to corresponding harsh environments of hot and cold deserts (e.g., Lee et al. 2011). While it is tempting to speculate that these duplications have been maintained through adaptation under selective pressure, we are aware that such ‘duplication’ events may simply be the remnants of ancient polyploidy or segmental duplication. In the absence of functional characterization, it is necessary to interpret all GO analyses with caution.

### Targeted analyses of drought and cold associated genes

We examined 29 functionally annotated genes identified to be involved in drought and cold tolerance (table 2), as several lineages have repeatedly colonized cold and dry environments. Most of these genes experienced significantly higher duplications than expected from WGDs including within Didiereoideae, Montiaceae and Cactaceae (figs. S6 and S7). Several of these genes may be good targets for future study. For example, WIN1 (SHN1) proteins, transcription factors associated with epicuticular wax biosynthesis that increase leaf surface wax content, showed duplications in both Didiereoideae and Montiaceae (table 2). Six homologs that are part of the calcium-dependent protein kinase (CDPK or CPK) gene family are known to be involved in drought stress regulation (Geiger et al. 2010; Brandt et al. 2012) and experienced several duplications: four at the origin of Didiereoideae, one within Montiaceae, one at Portulacaceae, and two within the Cactaceae.

**Table 2.** Selection analyses on 29 genes associated with cold and drought responses.

| Gene | TN | TS | Cac35 | Did27 | Mont15 | Port71 | SH80 (duplication) |
| --- | --- | --- | --- | --- | --- | --- | --- |
| **ABI1/PP2Cfamily** | 151 | 17 | 6 | 2 | 4 | 0 | 27 |
| NAC10 | 158 | 16 | 6 | 4 | 3 | 1 | 27,39 |
| WRKY33 | 165 | 13 | 4 | 2 | 3 | 1 | 17,27 |
| NAC29 | 163 | 13 | 7 | 2 | 2 | 0 | 13 |
| CDPK18 | 196 | 13 | 6 | 1 | 0 | 0 | 2,13,21,35,57 |
| **PP2C37 (PP2CA)** | 179 | 11 | 6 | 1 | 2 | 0 | 13,17,19,35 |
| HSP70 | 270 | 10 | 3 | 1 | 1 | 2 | 0,19,27,33,49,51,52 |
| ICE1 | 194 | 10 | 4 | 1 | 3 | 0 | 17,20,27,38 |
| HSP70 | 221 | 10 | 2 | 0 | 2 | 0 | 0,27,35 |
| ERF5 | 86 | 9 | 1 | 2 | 3 | 0 | 17,27,30 |
| HSP90-5 | 213 | 8 | 2 | 2 | 1 | 1 | 26,27,34 |
| CDPK1 | 85 | 7 | 4 | 2 | 1 | 0 | --- |
| SNRK2.5 | 156 | 5 | 2 | 1 | 0 | 1 | 39 |
| MYB124 (FLP) | 178 | 5 | 2 | 1 | 0 | 1 | 39 |
| HSP83 | 305 | 5 | 1 | 1 | 0 | 0 | 0,13,17,27,34,76 |
| CRPK1 | 177 | 5 | 3 | 2 | 0 | 0 | 13,20,27 |
| AREB1 | 92 | 4 | 1 | 1 | 1 | 0 | 17,27 |
| NAC2 | 157 | 4 | 2 | 0 | 0 | 1 | 27,39 |
| MYB60 | 87 | 4 | 2 | 0 | 0 | 1 | 27 |
| WIN1 (SHN1) | 79 | 4 | 0 | 0 | 0 | 1 | 17,27 |
| CDPK13 | 156 | 3 | 0 | 0 | 0 | 2 | 27 |
| CDPK26 | 117 | 3 | 1 | 1 | 0 | 1 | 0,17,27 |
| PLC2 | 108 | 3 | 1 | 0 | 1 | 0 | 17,19,35 |
| CDPK8 | 184 | 2 | 1 | 0 | 0 | 0 | 13,27,71 |
| CRLK1 | 170 | 2 | 2 | 0 | 0 | 0 | 27 |
| SNRK2.6 | 180 | 2 | 2 | 0 | 0 | 0 | 27,39 |
| SNRK2.4 | 125 | 1 | 0 | 0 | 0 | 0 | 17,27 |
| CDPK10 | 85 | 1 | 1 | 0 | 0 | 0 | 27 |
| ERF2 | 84 | 0 | 0 | 0 | 0 | 0 | 17,27 |

NOTE.—TN: tip number; TS: Total lineages under positive selection; Number of lineages under positive selection within certain clades, including Cactaceae (Cac35), Didiereoideae (Did27), Montiaceae (Mont15), and Portulacaceae (Port71). SH80: Duplication nodes with SH-like support > 80, Node number can be referred to in figs. 1 and S9. The bolded genes (ABI1 and PP2CA) are important negative regulators in the ABA pathway that mainly discussed in the text. More information about references and gene duplication patterns can be found in table S4.

These 29 genes also exhibited variable lineage-specific positive selection within Portulacineae (table 2). The strongest signals of positive selection were all found in genes associated with drought and/or cold tolerance in the ABA signaling pathway (such as the NAC10/29 and WRKY33 TFs, the CDPK18 and the PP2Cs, e.g., Golldack et al. 2014; Nakashima et al. 2014; Huang et al. 2015; Li et al. 2017), a central regulator of abiotic stress resistance. Interestingly, the ABI1 and PP2CA genes, which encode two proteins of the PP2Cs family, were among the genes with the highest number of lineages under positive selection. As negative regulators, PP2Cs can regulate many ABA responses, such as stomatal closure, osmotic water permeability, drought-induced resistance, seed germination, and cold acclimation (Gosti et al. 1999; Merlot et al. 2001; Mishra et al. 2006). In addition to PP2C genes, the positive regulators SNF1-related protein kinase 2s (SnRK2s) are also related to ABA signaling (Hubbard et al. 2010). While we did not detect many lineages under positive selection in the three SnRK2s genes (i.e., SnRK2.4, 2.5, 2.6, table 2), we found that they experienced ancient duplications at the origin of Portulacineae or earlier and recent duplications within Montiaceae, Didiereaceae, or Cactaceae (nodes 17, 27, 39; table 2). Positive selection and gene family expansion in these gene families within Cactaceae, Didieraceae, and Montiaceae suggests that these genes warrant further investigation for their potential role in the evolution of adaptations to challenging environmental conditions.

While several of these genes experienced duplications within clades of the Portulacinieae, many occurred at the origin of Portulacineae or earlier (table 2, table S4 and fig. S7). This suggests that the ability for the Caryophyllales, and the Portulacineae, to repeatedly adapt to harsh environments may have arisen early in the evolution of the clade and that the predisposition to become adapted to these environments may be the result of the early diversification of the gene families.

## Conclusions

WGDs (i.e., paleopolyploidy) have played critical roles in major events of plant evolution, but the nature and scale of their influence on macroevolutionary patterns and within individual clades is still debated. Our results suggest that WGDs in the Portulacineae were associated with several evolutionary patterns and processes. In addition to identifying significant gene-tree/species-tree discordance immediately after extensive gene duplication events, suggesting an association between rapid speciation and gene duplication and loss, we also found evidence for the association of WGDs and shifts into different environments and climates (Smith et al. 2018a). While this alone does not suggest specific cause, it suggests one means by which WGD can impact adaptation in plants and, perhaps, why some clades exhibit diversification shifts after WGDs. However, we did not detect association between WGD and diversification rate.

We also examined expanded gene families, GO over-representation, and selection within genes to determine whether potential molecular evolutionary patterns corresponded to adaptation to extreme environments. While these results provide essential resources for further and deeper examination with more detailed methods, they also highlight the major limitations in resources for non-model organisms. They rely on annotations based on model organisms that may be distantly related to the species of interest. While the gene itself may exhibit interesting patterns, the associated GO label may or may not carry much meaning and may have different functions in these different lineages. Despite these limitations, however, these analyses identified gene families that should be explored further, regardless of the accuracy of the GO label, as the patterns of selection and duplication alone suggest their importance.

Finally, our findings, contribute to a growing literature suggesting the importance of gene tree examination along with species tree construction (Smith SA et al. 2015; Brown and Thomson 2017; Shen et al. 2017; Walker et al. 2018b). Gene tree conflict is common and needs to be incorporated into our analyses of molecular and macro-evolution. The conflicts within these data likely hold interesting evolutionary questions and answers that should be analyzed in more detail.

## Material and Methods

### Taxon sampling, transcriptome generation and homology inference

Sixty-eight ingroup species were included in this study, representing all families within the Portulacineae (Anacampserotaceae, Cactaceae, Basellaceae, Didiereaceae, Montiaceae, Talinaceae, and Portulacaceae) *sensu* APG IV (Angiosperm Phylogeny Group, 2016) except for Halophytaceae. We also included 14 Caryophyllales species as outgroups, including seven taxa of Molluginaceae (tables S5 and S6). Of the 82 transcriptomes included in the study, 47 were newly generated following the protocol of Yang et al. (2017). We followed the pipeline of Yang and Smith (2014) with minor modifications (available at https://bitbucket.org/ningwang83/Portulacineae) to infer homologs. In total, we obtained 8,592 final homolog clusters, which are used in orthology inference, gene duplication analyses and GO annotation. More details about transcriptome generation and homology inference can be found in Supplementary Material and Methods.

### Orthology Inference and Species Tree Estimation

We used the rooted tree (RT) method in Yang and Smith (2014) to extract orthologs from homolog trees with *Beta vulgaris* as the outgroup. Orthologous clades with more than 30 ingroup taxa were retained (fig. S8). We extracted orthologs with > 75 taxa, aligned and cleaned them with MAFFT v7 (genafpair –maxiterate 1000, Katoh and Standley 2013) and Phyutility (-clean 0.3, Smith and Dunn 2008), and inferred a ML tree for each using RAxML v8.1.22 (Stamatakis 2015) with GTR+Γ model. After removing taxa with a terminal branch length longer than 0.1 and 10 times greater than sister clade from 58 orthologs, 841 orthologs including at least 77 taxa and 500 aligned DNA characters were re-aligned with PRANK v.140110 (Löytynoja and Goldman 2008) using default settings and trimmed in Phyutility (-clean 0.3).

The species tree was inferred using two methods. First, a concatenated matrix was built with the 841 orthologs and a ML tree was estimated by using RAxML with the GTR+Γ model partitioned by gene. Node support was evaluated by 200 fast bootstrap replicates. Second, an MQSST tree was estimated in ASTRAL 4.10.12 (Mirarab et al. 2014) using ML gene trees, with uncertainty evaluated by 200 bootstrap replicates using a two-stage multilocus bootstrap strategy (Seo 2008).

### Assessing conflicts among gene trees

We assessed conflict by mapping the 841 rooted ortholog trees onto the species tree topology with Phyparts (Smith SA et al. 2015). Trees were rooted on *Beta vulgaris* if present, using *Limeum aethiopicum* and/or *Stegnosperma halimifolium* if *Beta* was not present, and with *Sesuvium portulacastrum*, *Delosperma echinatum*, *Anisomeria littoralis* and *Guapira obtusata,* if the first three species were not present. We only considered nodes with >= 70% bootstrap support in the gene trees. We conducted likelihood comparisons in more detail as in xWalker et al. (2018b) for three areas of the tree: 1) Cactaceae and relatives, 2) the early diverging branch within Cactaceae (i.e., *Leuenbergeria* clade), and 3) with the placement of Basellaceae and Didiereaceae. We calculated likelihood scores for each gene tree constrained to alternative resolutions and compared scores. We considered a higher ln *L* for one topology as a sign of support, and |Δln *L|* (the difference in ln *L* value) > 2 as a sign of significant support (as in Walker et al. 2018b). Potential outlying genes that exhibited extreme deviations in likelihood were examined for lineage specific positive selection. We analyzed potential hybridization with PhyloNet (Than et al. 2008) and D_FOIL_ (Pease and Hahn 2015). PhyloNet does not run well on large datasets so we pruned the gene trees to include 5 taxa from Cactaceae, one from each ingroup family, with the *Limeum aethiopicum* as an outgroup. Species were selected based on their gene occupancy. We conducted Maximum Pseudo Likelihood analyses with 10 maximum reticulations. D_FOIL_ analyses can only analyze 5-taxon trees, so we reduced gene trees to four species with the highest gene occupancy for the clade of interest and *Pharnaceum exiguum* as an outgroup. For example, we choose two species from Cactaceae and one from each of Portulacaceae and Anacampserotaceae, to test the controversial relationships among these families.

### Inference of gene and genome duplication

To infer gene duplication, we considered nodes in gene trees with SH-like support > 80. We extracted 8,332 rooted clusters with >= 30 taxa, and conducted Phyparts analyses. We also conducted Ks plot analyses following the same process of Yang et al. (2015). Briefly, we reduced highly similar PEP sequences (by CD-HIT: -c 0.99 -n 5, Fu et al. 2012) for each of the 82 species, and conducted an all-by-all BLASTP (-evalue = 10, -max_target_seq = 20) and removed highly divergent hits with < 20% similarity (pident) or < 50 aligned amino acids (nident). Sequences with ten or more hits were also removed to eliminate large gene families. We then used the PEP and corresponding CDS from paralogous pairs to calculate Ks values using the pipeline https://github.com/tanghaibao/biopipeline/tree/master/synonymous_calculation. Peaks were identified by eye as in Yang et al. (2015).

To determine whether a potential WGD event occurred before or after a speciation event, we calculated between-species Ks distribution using orthologous gene pairs. The procedure is similar as above, except that a reciprocal BLASTP was carried out between two species instead of an all-by-all BLASTP within one taxon. Between-species Ks plots were compared to within-species Ks plots to determine the relative timing of the WGD and speciation event.

### Reconstruction of climate occupancy and diversification rate

To test whether WGD were associated with climate niche shift and/or diversification rate shift, we reconstructed ancestral climate occupancy and diversification on a species level phylogeny of Portulacineae. First, sequence data from NCBI were gathered using PyPHLAWD (Smith and Brown 2018). To increase the accuracy of phylogenetic inferences, major clades were constrained based on the transcriptomic results here and were conducted on Molluginaceae and Cactineae (i.e., Portulacineae) individually and combined. ML and 100 bootstrap trees were constructed with RAxML. For divergence time estimation, we selected five genes from the transcriptomic dataset using SortaDate (Smith et al. 2018b). A time tree, hereafter ‘small tree’, was then built with them in BEAST 1.8.3 using 13 secondary time calibrations (table S7) from Arakaki et al. (2011). The species-level tree, hereafter ‘big tree’, as well as the 93 bootstrap trees that were consistent among family level relationships in the ‘small tree’ were then dated in treePL (Smith and O’Meara 2012) using the dates from the ‘small tree’ as constraints.

Reconstruction of ancestral climate occupancy state and diversification rate followed the methods of Smith et al. (2018a). Briefly, we extracted the climate occupancy data (COD) for the Portulacineae and Molluginaceae species from GBIF based on the taxa from the ‘big tree’. Mean climate values of Bioclim 1 (mean annual temperature), 12 (mean annual precipitation), and principal component axis 1 (based on a PCA of the set of full bioclimatic variables) were calculated for each taxon. Both ancestral states and Brownian motion rates of evolution were compared between WGD nodes and their sister clades. We reconstructed diversification using MEDUSA (Alfaro et al., 2009; Pennell et al., 2014) on the ‘small tree’ with the estimated number of species for each taxa group and the ‘big tree’.

### Annotation of expanded gene families

To investigate gene function and GO (gene ontology) term overrepresentation, we conducted several analyses. We conducted BLASTX analyses against the nonredundant protein NCBI database by using a PEP sequence from each of the top 20 genes with the highest total number of tips. We also identified clades with large number of gene duplications [i.e., > 160 (2%) gene duplications], and conducted BLASTN analyses (-evalue 10) of these against the database of *A. thaliana* (release-34 EnsemblPlants). GO terms and IDs were recorded. We conducted GO overrepresentation test (i.e., overrepresentation of genes in a specific functional category, http://www.pantherdb.org) using genes duplicated at the origin of Portulacineae (Node 13) as background to detect potential overrepresentation of genes that duplicated more recently. We corrected for multiple tests using a Bonferroni correction.

### Identification of lineage-specific selection on stress response genes

We carried out selection analyses on a targeted set of 29 genes that were present in our data and are known to be associated with cold and/or drought adaptation. We repeated the homolog filtering as indicated above three times to reduce potential assembly or clustering errors. CDS sequences were aligned based on the corresponding peptide alignments using phyx (Brown et al. 2017). We further cleaned CDS alignments by combining sequences with identical overlapping segments to eliminate the potential assemble redundancy, which, although uncommon, may cause error in these analyses. After trimming the CDS alignment (Phyutility-clean 0.1), a ML tree was built for each homolog in RAxML with GTR+Γ model and rooted with outgroup species. Lineage specific selection was calculated for each homologous gene in HyPhy v2.2.4 (Pond and Muse 2005) using an adaptive branch-site random effects likelihood test (aBSREL, Smith MD et al. 2015), which automatically infers an appropriate model among branches. The optimal number of ω categories was selected according to AICc scores. The branches with episodic positive selection that show significant proportion of sites with ω > 1 were chosen with p < 0.05 after applying the Holm-Bonferroni multiple testing correction.

To test whether the 29 targeted genes associated with cold/drought adaptation experienced significant more duplications at certain nodes, we randomly selected 29 genes from the final homologous clusters (with an additional overlap checking to be compatible with the treatment of the targeted 29 genes) and calculate the number of genes duplicated. We repeated this process for 1000 times and plotted the frequency distribution of the number of duplicated genes for each node (fig. S7).

## Supplementary Material

Supplementary data are available at Molecular Biology and Evolution online.

## Acknowledgments

The authors thank Wynn Anderson, the Bureau of Land Management, the US Forest Service, and the staff of Desert Botanical Garden, Sukkulenten-Sammlung Zürich, Cambridge University Botanic Garden, Missouri Botanical Garden, and the Oberlin College Greenhouse for permission to collect specimens, and thank Hilda Flores, Helga Ochoterena, and Norman Douglas for help with collecting. The authors also thank Jeet Sukumaran for constructive discussion on gene tree discordance. Special thanks to Oscar Vargas for a thorough review of the paper. This work was supported by NSF DEB 1354048 to SAS and NSF DEB 1352907 to MJM.

